# 3D-Mass Spectrometry Imaging of Micro-scale 3D Cell Culture Models in Cancer Research

**DOI:** 10.1101/2022.12.05.519157

**Authors:** Stefania-Alexandra Iakab, Florian Keller, Stefan Schmidt, Jonas Cordes, Qiuqin Zhou, James L. Cairns, Frank Fischer, Richard Schneider, Ivo Wolf, Rüdiger Rudolf, Carsten Hopf

**Affiliations:** Center for Mass Spectrometry and Optical Spectroscopy (CeMOS), Mannheim University of Applied Sciences, Paul-Wittsack-Str. 10, 68163 Mannheim, Germany; Faculty of Computer Science, Mannheim University of Applied Sciences, 68163 Mannheim, Germany; Medical Faculty Mannheim, Heidelberg University, Theodor-Kutzer-Ufer 1-3, 68167 Mannheim; Merck KGaA, Frankfurter Str. 250, 64293 Darmstadt; Mannheim Center for Translational Neuroscience (MCTN), Heidelberg University, Theodor-Kutzer Ufer 1-3, 68167 Mannheim; Medical Faculty, Heidelberg University, Im Neuenheimer Feld 672, 69120 Heidelberg

**Keywords:** 3D cell culture imaging, colon cancer model, MALDI imaging, metabolomics, elastic 3D reconstruction

## Abstract

Three-dimensional (3D) human cell culture models have emerged as a key technology for personalized medicine and for phenotypic compound screening in more disease-like *in-vitro* systems. Mass spectrometry imaging (MSI) is one of the most versatile label-free techniques that enables simultaneous generation of spatial maps for multiple relevant molecules in these 3D-models. Here, we present an integrated platform for 3D-MSI of 3D-cell cultures comprising 3D-printed metal casting molds for freezing and embedding, MS imaging of 100 serial cryosections and their computational elastic 3D-reconstruction. With this platform, we monitored multiple lipids that were selectively associated with different cell-types or cell-cell interactions within 300 μm-scale fibroblast and colon cancer biculture spheroids. Our findings suggest that 3D-printing-aided precise preparation of serial sections from small spheroids and visualization of marker molecules in 3D can provide a detailed overview of the cellular metabolic interplay in 3D cell culture models in cancer research and drug discovery.

## Introduction

Three-dimensional (3D) human cell culture models are providing a diversified platform that is not only time- and cost-efficient, but also presents human rather than mouse enzymes and drug targets, is thought to mimic *in vivo* conditions and thus supports animal-free testing and search for personalized therapies. Pharmaceutical research and development is highly interested in such models for early assessment of drug dosage, drug formulations, in-tissue localization and biological responses[1]. Therefore, while 3D cell cultures are already critical resources in pre-clinical screening, their routine deployment for use with high molecular content analytical methods such as matrix-assisted laser desorption/ionization (MALDI) mass spectrometry imaging (MSI) is still lacking and has become an important goal.

In the context of malignant disorders, such as colorectal cancer, the metabolic interplay between cancer and stromal cells has become an interventional target. Indeed, aerobic glycolysis[2] and mutual exchange of metabolites[3] have been identified as important components of the malignant phenotype. To model the metabolic relationship between malignant and stromal cells, Keller et al. developed colon cancer spheroids composed of human HT-29 colon cancer cells and human CCD-1137Sk fibroblasts[4]. This provided evidence for the occurrence of reverse Warburg effect[5], where stroma cells exhibit reduced oxidative metabolism and deliver energy-rich metabolites to adjacent malignant cells. In this example and others, to further increase the mechanistic understanding of the metabolic interplay between malignant and stroma cells, the spatially resolved detection of metabolites and lipids would be highly beneficial.

MALDI-MSI is a label-free technology that can provide fast, accurate and comprehensive molecular information. It is equally useful in targeted, quantitative MSI of known drugs or metabolite markers and in untargeted analysis in marker discovery and drug safety assessment[6–9]. MSI was first applied to 3D cell culture models in 2011 to map protein distribution within a colon carcinoma model[10]. Since then, several studies have used MALDI-MSI to characterize 3D cell culture models with both targeted and untargeted strategies. For example, breast cancer-associated metabolites were identified within regions with low or high oxygen availability inside spheroids[11], drug penetration profiles of the PI3K/AKT/mTOR pathway disrupting alkyl-phospholipid perifosine were examined in colorectal carcinoma spheroids[12], and the metabolomic and lipidomic response to the chemotherapeutic drug doxorubicin was investigated in an osteosarcoma spheroid model[13]. Most of these studies focused on analyzing isolated 2D sections of complex samples, thereby neglecting the 3-dimensional nature and a major advantage of spheroids: their small size. Indeed, these samples can be measured rapidly, thus allowing scans of multiple consecutive sections for 3D-reconstructions of the entire spheroid in space; along these lines, also size of such data sets is sufficiently small to allow 3D data processing and reconstruction within the same software[14]. Based on molecular patterns, three principal regions of spheroids, i.e., necrotic core, annular quiescent region, and proliferative outer region, were suggested in 3D[15]. Thus, although MSI enables exploring multiple molecular distributions of a spheroid in a label-free manner and although it is not limited to the analysis of protein distribution, there is no integrated platform yet for spheroid MSI sample preparation with the precision needed for subsequent 3D-reconstruction. Indeed, major technical hurdles that all result in poor reproducibility include: i) the very small sample size, which makes handling and visual detection during cryo-sectioning very challenging, ii) sample damage during sectioning[16], and iii) the difficulty of collecting and mounting unperturbed consecutive sections of sufficient quality for 3D-reconstruction. One initial protocols for step-by-step sample preparation[17] has been followed, with some adaptations, by other groups[11, 12]. Other studies employed different approaches, e.g., by using commercial negative molds[18], in-house built gelatin blocks[13], or different embedding media[3, 15].

To address the technical hurdles mentioned above, here, we introduce a platform that combines the following elements: We designed a 3D-printed metal casting mold for reproducible generation of two-component gelatin cryo-molds and conceptualized a pertinent workflow for optimal sample preparation of spheroids for 3D-MSI. This enabled reliable collection of over 40 consecutive sections from stacks of 300 μm spheroids. Reliable sampling of entire spheroids allowed to monitor lipids/metabolites associated to different cell-types or cell-cell interactions within a colon cancer-fibroblast co-culture spheroid model in 3D. Moreover, the high quality of consecutive sections enabled subsequent computational elastic reconstruction of the three-dimensional distribution of specific molecular species employing open-source M^2^aia software[19]. Analysis of reconstructed spheroids demonstrated (cell-type-) specific heterogeneous enrichment of several molecular entities in space, demonstrating the advantage and relevance of the approach. Owing to the inherent technical reproducibility, our approach also increases the throughput of MSI analysis of spheroids, which could lay the foundation for 3D-MSI based drug profiling.

## Materials and methods

### Chemicals

Polyvinylpyrrolidone (PVP) (MW 360 kDa), (Hydroxypropyl)-methylcellulose (HPMC) (viscosity 40-60 cP, 2 % in H_2_O (20 °C), N-(1-naphthyl) ethylenediamine dihydrochloride (NEDC)[20], 1,5-diaminonaphthalene (DAN), 1,1’-Binaphthyl-2,2’-diamine (BNDM)[21], trifluoroacetic acid (TFA), 9-aminoacridine (9-AA) and ethanol were purchased from Sigma Aldrich (Taufkirchen, Germany); 2,5-dihydroxybenzoic acid (DHB) was purchased from Alfa Aesar (Karlsruhe, Germany), 4-Phenyl-α-cyanocinnamic acid amide (PhCCAA)[22] from SiChem (Bremen, Germany), dimethyl sulfoxide (DMSO) from VWR (Darmstadt, Germany), carboxymethyl cellulose (CMC) from Fluka (VWR, Darmstadt, Germany), gelatin 180 bloom from Carl Roth (Karlsruhe, Germany), methanol and acetonitrile (ACN) from Honeywell (VWR, Darmstadt, Germany), and ESI tune mix form Agilent Technologies (Santa Clara, United States). All solvents used were of analytical grade or higher.

### 3D cell culture models

Cells were cultured following the protocol from Keller et al[4]. In brief, CCD-1137Sk human fibroblasts purchased from ATCC (LGC Standards GmbH, Wesel, Germany) were kept in Iscove’s Modified Dulbecco’s Medium (Capricorn; IMDM-A; Lot #CP22-5149) supplemented with 10% fetal bovine serum (Capricorn; FBS-12A; Lot #CP20-3380), and 1 % penicillin/streptomycin (Capricorn; PS-B; Lot #CP21-4079). HT-29 human colon cancer cells (purchased from ATCC, LGC Standards GmbH, Wesel, Germany) were cultivated in McCoy’s 5a media (Capricorn; MCC-A; Lot #CP22-5172), supplemented with 10 % FBS-12A and 1 % PS-B. Cell lines were passaged twice per week with a seeding density of 1 × 10^6^ cells/T75 flask for HT-29 cells and 1.5 × 10^6^ cells/T75 for CCD-1137Sk. Mono-cultured 3D models (or spheroids) were obtained using a total of 10 000 cells seeded per well for each cell line and the co-cultured spheroids were obtained by seeding 10 000 cancer cells together with 10 000 fibroblasts, using 96-well cell-repellent microplates (faCellitate; BIOFLOAT; F202003) that were centrifuged (7 min / 500 rcf) prior to further cultivation.

### Embedding and cryo-sectioning of spheroids

Spheroids were embedded with the help of a gelatin cryo-mold created using a 3D printed metal casting mold developed in-house. The metal base was directly printed as a positive module using austenitic steel powder (MetcoAdd 316L-A, Oerlikon Metco Europe GmbH, Switzerland) with a 3D metal printer (TruPrint 2000, Trumpf, Germany). Autodesk Inventor Professional 2022 (Autodesk Inc., USA) was used to compile the CAD designs. For creating the gelatin cryo-mold, approximately 3 ml of 350 mg/ml gelatin solution (in water) was poured into the metal casting mold and left to solidify at −20 °C for 15 minutes. The gelatin block was removed and used immediately. 7.5 % HPMC-2.5 % PVP solution was pipetted into the channels of the gelatin cryo-mold at room temperature. Several spheroids were harvested in a 2 ml Eppendorf tube, and then transferred directly into the HPMC-PVP-filled channels using a pipette. Finally, the gelatin/HPMV-PVP block filled with spheroids was immediately snap-frozen in liquid nitrogen. Cryo-sectioning was performed on a CM1950 cryostat (Leica Biosystems GmbH, Nussloch, Germany). Frozen gelatin blocks were mounted with CMC on the sample holder and kept inside the cryostat chamber (temperature set to −20 °C) for 15 min prior to sectioning. Spheroid sections were mounted on conductive ITO coated slides (Diamond Coatings Ltd, West Midlands, UK) or Intellislides (Bruker Daltonics, Bremen, Germany) for MALDI-MSI. For 2D imaging, 10 μm thick sections were used, and for 3D reconstruction more than 50 consecutive sections were collected on one or two ITO slides. Slides were either used right after sectioning or stored at −80 °C. All slides were desiccated at low pressure for ~15 min before further MALDI-MSI sample preparation.

### MALDI mass spectrometry imaging

Five matrices were evaluated for MALDI-MSI of spheroids: PhCCAA (3 mg/ml in 70 % ACN), 9AA (6 mg/ml in 70 % ACN), DAN (6 mg/ml in 70 % ACN), NEDC (10 mg/ml in 70 % methanol), BNDM (10 mg/ml in 70 % ACN) and DHB (30 mg/ml in 70 % ACN). The matrices were sprayed using an HTX-TM3 sprayer. The spraying parameters are indicated in Supplementary Table S1. Data was acquired on a timsTOFfleX mass spectrometer (Bruker Daltonik, Bremen, Germany), operated in negative ionization mode, in tims OFF mode. The mass range was between *m/z* 50 and 1200 with a spatial step-size of 20 μm, and between 50-200 laser shots were summed up per pixel. For all experiments, the laser was operated with a repetition rate of 10 kHz, with some exceptions at 5 kHz. All raw data was directly uploaded and processed in SCiLS lab (Version 2023a Pro). All data shown (ion images and spectra) were normalized to the root mean square (RMS). Feature annotation was done by exact mass (<10 ppm mass error).

### Computational 3D-MSI reconstruction and visualization

The 3D reconstruction was done using M^2^aia [19] v2022.08 (biotools:m2aia), an open-source software dedicated to memory-efficient and fast MSI data visualization, to simultaneous handling and registration of multiple, potentially multi-modal images, and to 3D MS image reconstruction. Data collected from consecutive sections was previously RMS normalized and then imported as individual centroid imzML files, then the morphology-representative ion *m/z* 863.5 was chosen for the automatic stacking procedure. Default parameters were used with a maximum of 20 rigid iterations and 400 deformable iterations. 3D stacks were then exported in .nrrd file format and imported into Slicer 3D (https://www.slicer.org/), an open source software for creating the 3D model. Blender open source software (https://www.blender.org/) was used for obtaining volume visualization videos.

## Results and Discussion

### 3D-metal printing-assisted sample preparation workflow for spheroids

3D cell culture models are an attractive technology for pre-clinical compound screening. However, sample preparation for 3D-reconstruction following MALDI-MSI analysis is not trivial. In most cases, the size of the cultures is in the range of few hundreds of micrometers, which is challenging for the handling and requires very precise and gentle procedures. Apart from the dexterity needed in the laboratory, pioneering spheroid sample preparation methods[17] mount a relatively small number of sections per slide. This may introduce unnecessary batch effects between measurements and requires more MALDI sample preparation materials, time, and effort. To overcome these key hurdles in 3D-MSI of 3D-cell cultures, we first designed 3D-printed metal casting molds (with three or nine channels, **Figure S1**) as support for casting of gelatin cryo-blocks for the embedding of several 3D-culture samples in a precise and reliable sterical arrangement (**Figure S2**). **Figure 1** illustrates the full workflow, from harvesting of cultured spheroids to embedding them in HPMC-PVP-filled gelatin cryo-molds, and to finally freezing them in liquid nitrogen for cryo-sectioning or storage. Our method requires only standard laboratory equipment and diligence from the user when transferring the spheroids into the embedding medium. It offers reproducible and easy sample preparation, enhanced throughput (when using the nine-channel gelatin cryo-mold), and, most importantly for 3D-reconstructions, reliable consecutive sectioning (**Figure S3**).

**Figure 1.**
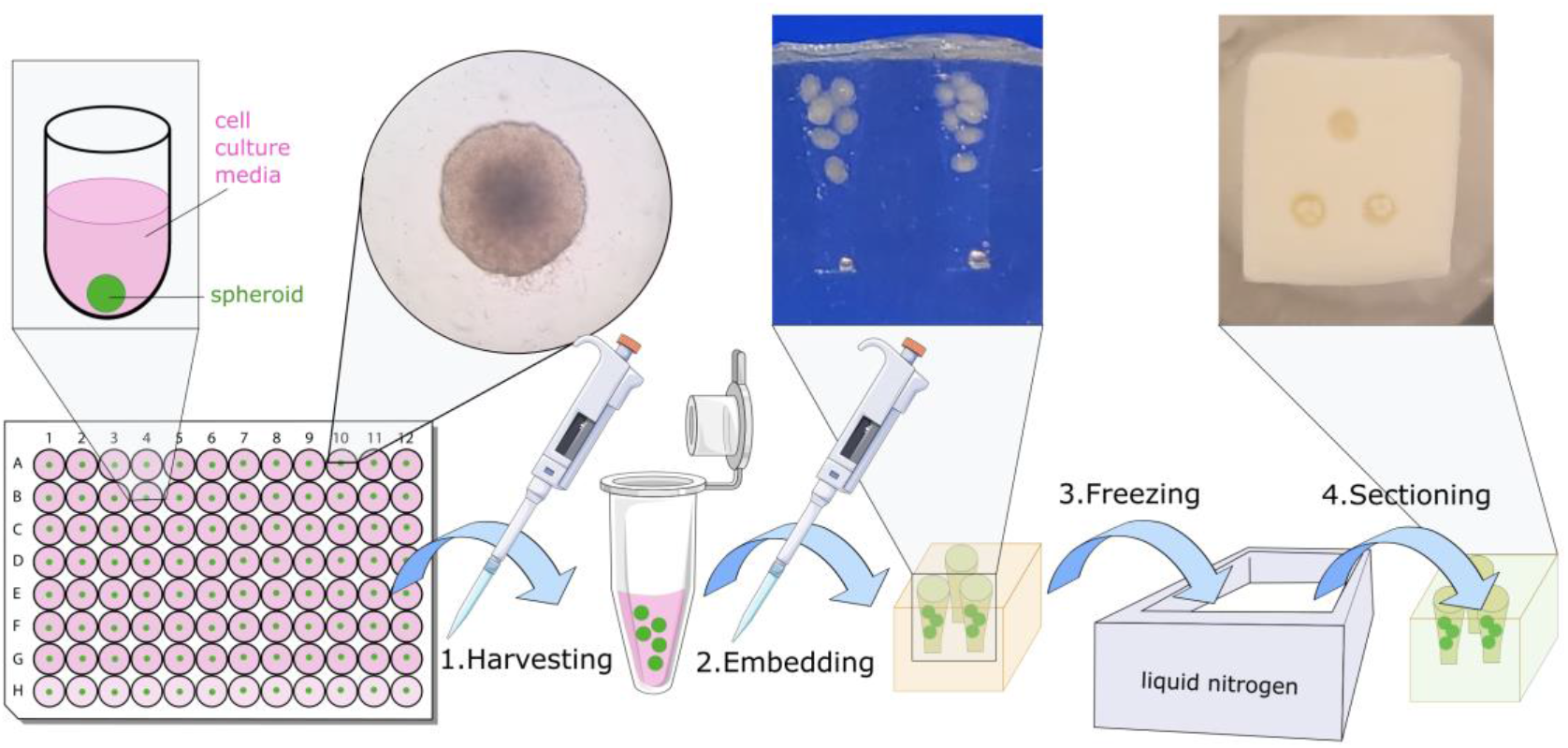
Sample preparation protocol. Spheroids grown in a 96-well plate are harvested, then embedded in HPMC-PVP-filled gelatin cryo-molds, snap frozen in liquid nitrogen, and finally cryo-sectioned. Inserts showcase pictures from the CeMOS laboratory.

### MALDI matrix selection for spatial metabolomics in serial cancer spheroid sections

To optimize MALDI MS imaging specifically for spatial metabolomics in serial spheroid sections, we optimized a key aspect of MALDI-MSI sample preparation, namely matrix selection. The co-culture spheroid model was designed to investigate the metabolic interplay between cancer and stromal cells[4], the so-called reverse Warburg effect that involves metabolic shuttling of metabolites like lactate, pyruvate and tricarboxylic acid (TCA) cycle metabolites. As metabolites of glycolysis and TCA cycle can be detected as negative ions in MALDI-MSI[15], we chose six MALDI matrices that are commonly used for detecting small molecules (mass range 100-1200 Da) in negative ionization mode. We evaluated the performance of PhCCAA, 9AA, DAN, NEDC, DHB, and BNDM by comparing their sensitivity (coverage of the mass spectra and peak intensity) and specificity (number of *m/z* features detected only from the spheroids).

The mean spectrum collected from the spheroids spray-coated with the respective matrices is illustrated in **Figure 2**. In all cases, the higher mass region showed a rich lipid peak distribution, while in the lower mass range, in some cases (DAN, 9AA, DHB), the matrix peaks were the most intense. To better understand the sensitivity of each matrix, we generated an intensity heatmap for a list of most abundant spheroid-specific features which were selected following a ROC analysis between matrix and spheroid pixels (**Figure 3A**). NEDC displayed the widest detection coverage, although its sensitivity for most features was comparable to DAN and BNDM. The ion images of a low-abundant small molecule (*m/z* 303.23, annotated as the [M-H]^-^ form of arachidonic acid) and a highly abundant lipid (*m/z* 863.56, annotated as the [M-H]^-^ form of PI 36:1) indicated that NEDC yielded the most clear-cut ion distribution on the spheroid surface, while DAN and BNDM showed slight diffusion or delocalization of the molecules (**Figure 3 B and D**). To avoid user-biased evaluation, we also introduced a sensitivity threshold, based on the average intensity per image. In our case, this threshold was 100 arb. units, which was chosen based on the peak filter threshold used in SCiLSlab software to select the spheroid-specific features. All results supported NEDC as the most suitable matrix for detecting small molecules in negative mode from these spheroids.

**Figure 2.**
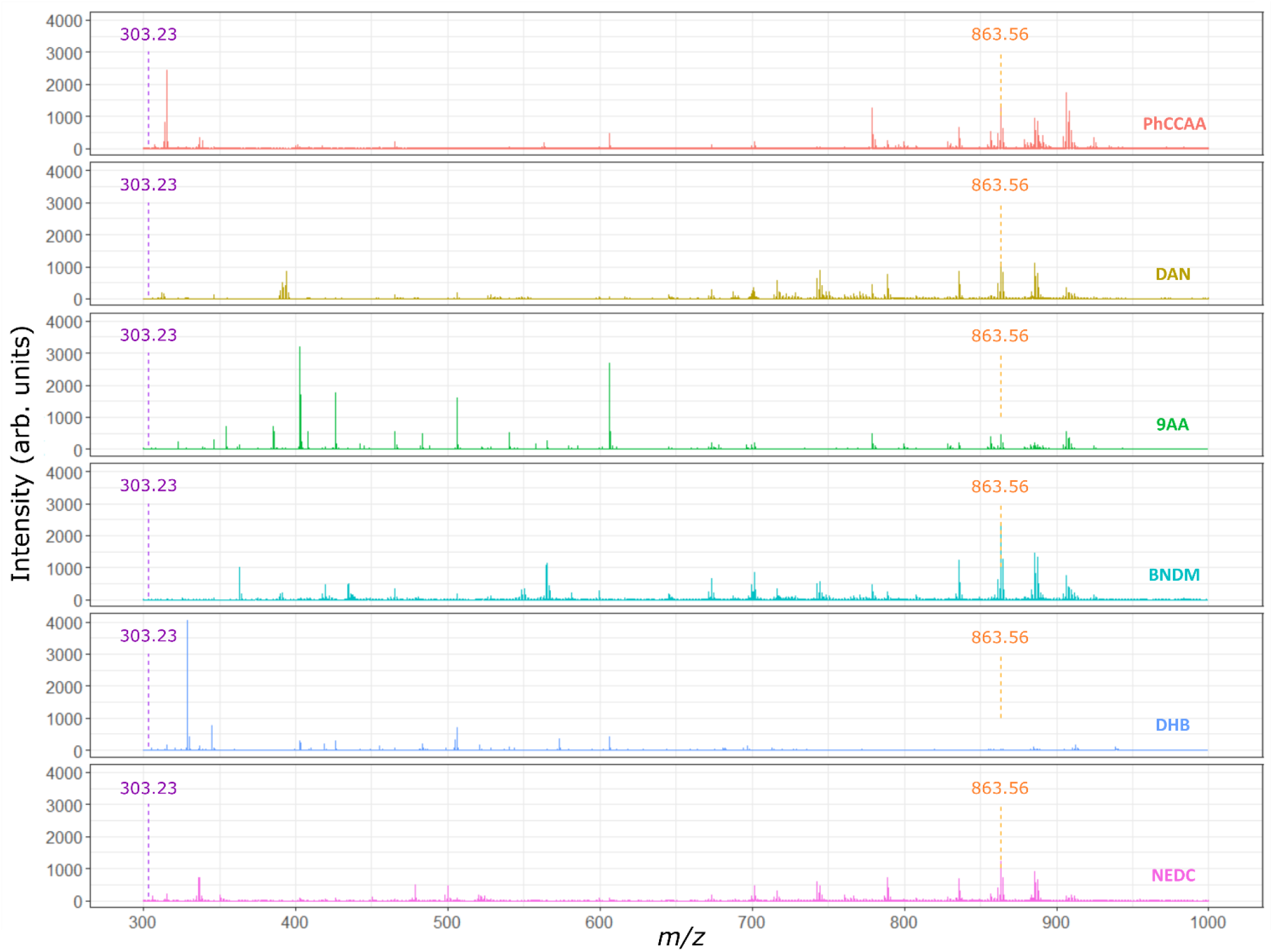
Mean spectra collected from spheroids sprayed with different matrices. MALDI-MSI mean spectrum from spheroid-only regions measured using PhCCAA (red), DAN (yellow), 9AA (green), BNDM (light blue), DHB (dark blue) and NEDC (pink). Data were RMS normalized. Highlighted with dashed lines are the low abundance metabolite *m/z* 303.2328 (in purple), and a high abundance lipid at *m/z* 863.5687 (in orange).

**Figure 3.**
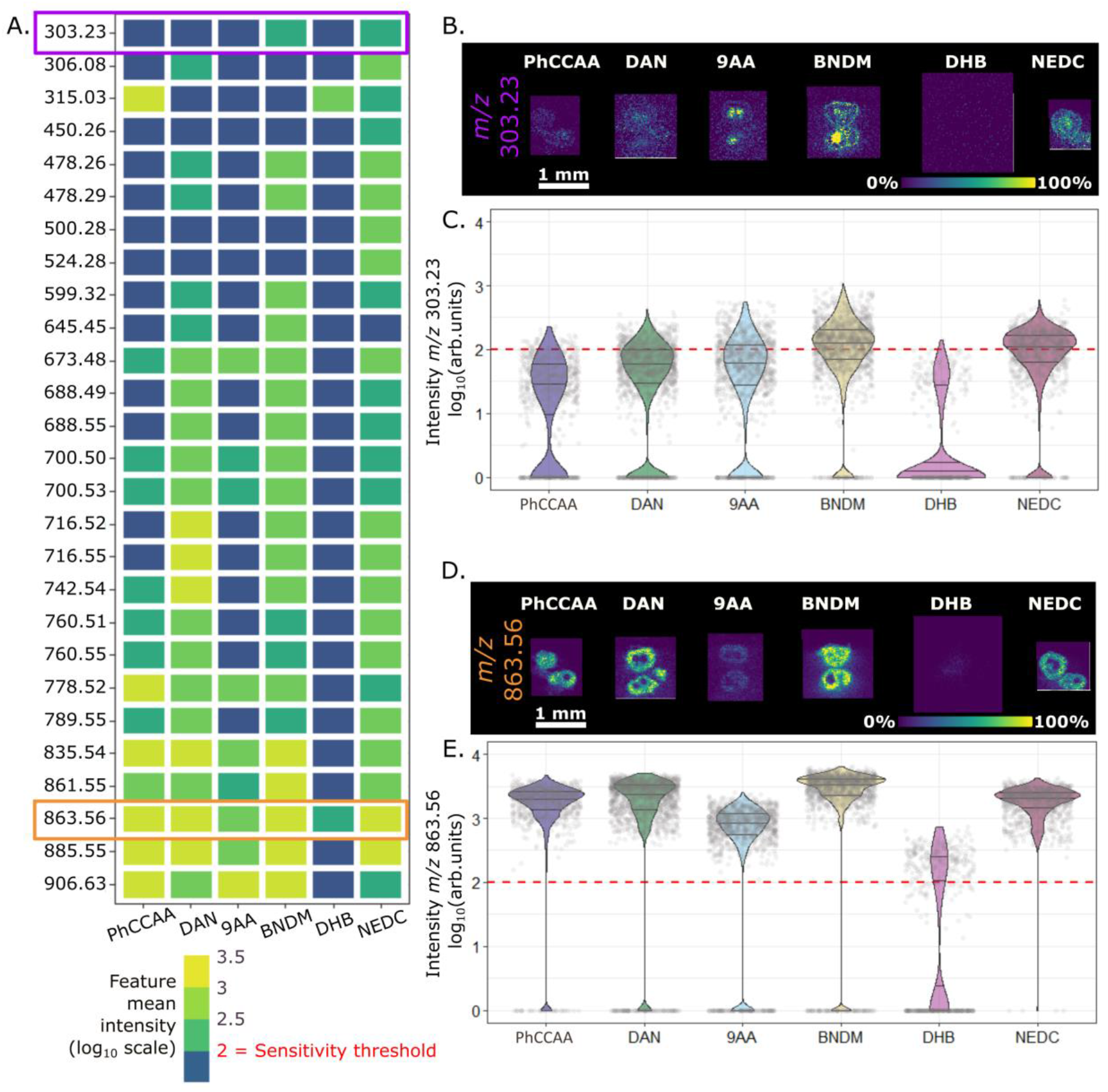
Matrix selection. Heatmap of spheroid-specific features detected by 6 matrices (A); the sensitivity threshold was set to 100 a. u. based on the peak filter threshold used in SCiLS; mean intensity values were obtained only from spheroid-specific pixels (ROIs obtained by bisecting k-means clustering in SCiLS using the list of features from the full data set, by setting the filter threshold at 360 (absolute intensity value)). Highlighted in purple is the low abundance metabolite *m/z* 303.23, and in orange, the high abundance lipid at *m/z* 863.56. Ion images from each matrix of *m/z* 303.23 (B) and *m/z* 863.56 (D) and the associated violin plots for each image (C) and (E), respectively.

### Elastic 3D-MS imaging reconstruction of spheroids and data analysis

Small molecules are the essence of cancer metabolism, but there are few analysis methods that can give spatial distribution pictures of a wide range of molecules simultaneously or to analyze complex metabolite patterns with spatial resolution. We used two MALDI-MSI methods to obtain high quality imaging data in the ranges of small molecules (*m/z* 50-600 Da, shown in supporting information) and lipids (*m/z* 300-1200 Da), from which we were able to detect over 200 spheroid-associated molecules. First, the data collected with the timsTOFfleX were imported to SCiLSlab software, where an *m/z* feature list was generated by setting an intensity threshold to 300 a.u. Using this raw feature list, we then segmented the data cubes into two clusters, which separated spheroid from background or matrix pixels. The root-mean-squared (RMS) normalized average spectrum collected from the spheroid is shown in **Figure 4A**, along with spheroid-specific features presumed to be lipid species labelled in **Figure 4B**.

**Figure 4.**
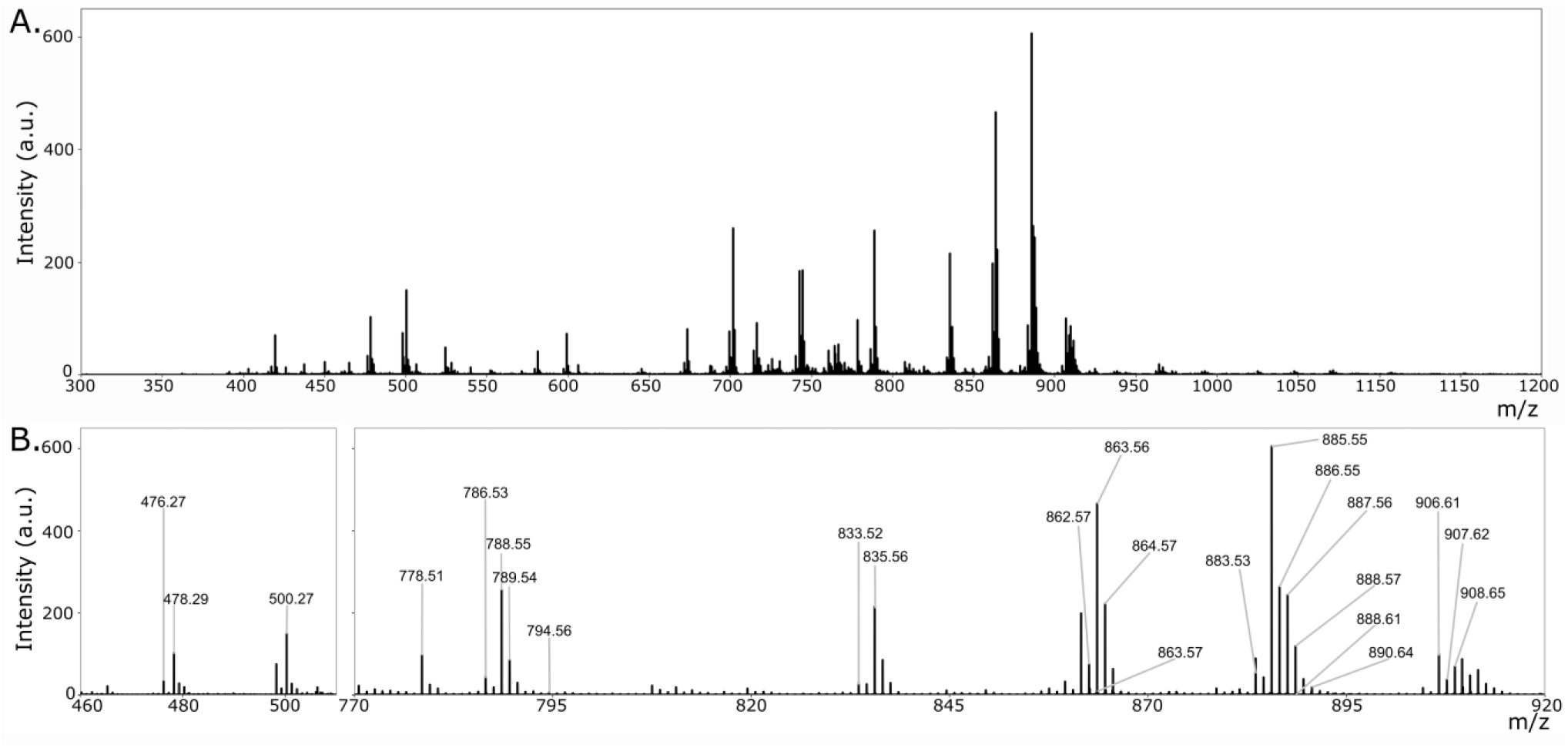
Putative lipid species detected from the spheroid model. Mean spectrum from the spheroid specific pixels using NEDC MALDI matrix in negative ion mode covering the full m/z-range (**A**). Highlighted regions from the mean spectrum with *m/z*-values of putative lipid species (**B**).

We next proceeded with unsupervised multivariate data analysis, i.e., segmentation and principal component analysis (PCA). Interestingly, multiple different spectral patterns appeared, two presumably corresponding to fibroblasts in the core and cancer cells outside the core, and two which correspond to more than one molecular composition within a given cell-type specific position. Indeed, the segmentation images (**Figure 5A**) – obtained using bisecting *k*-means clustering – revealed four distinct regions within the bi-cultured spheroids: (i) a core region (in green), (ii) a spatially restricted “communication” region (in yellow), (iii) a wider outer region (in purple) and (iv) an external margin region (in orange). Referring to our published study[4], these likely corresponded to (i) the fibroblast core, (ii) the contact zone between fibroblasts and cancer cells, (iii) the cancer cells, and (iv) the outer cancer cell ring. This prompted us to investigate possible cell type-specific and cell-cell interaction biomarkers. PCA supported this interpretation. Indeed, while pixels assigned to fibroblasts and the communication layer grouped together in PC2 and were separated by PC1, the pixels from the cancer cells and the exterior layer grouped together in PC1 and were separated by PC2 (**Figure 5BC**). This suggested that the positioning of a specific cell type within a spheroid changed that cell types molecular composition.

**Figure 5.**
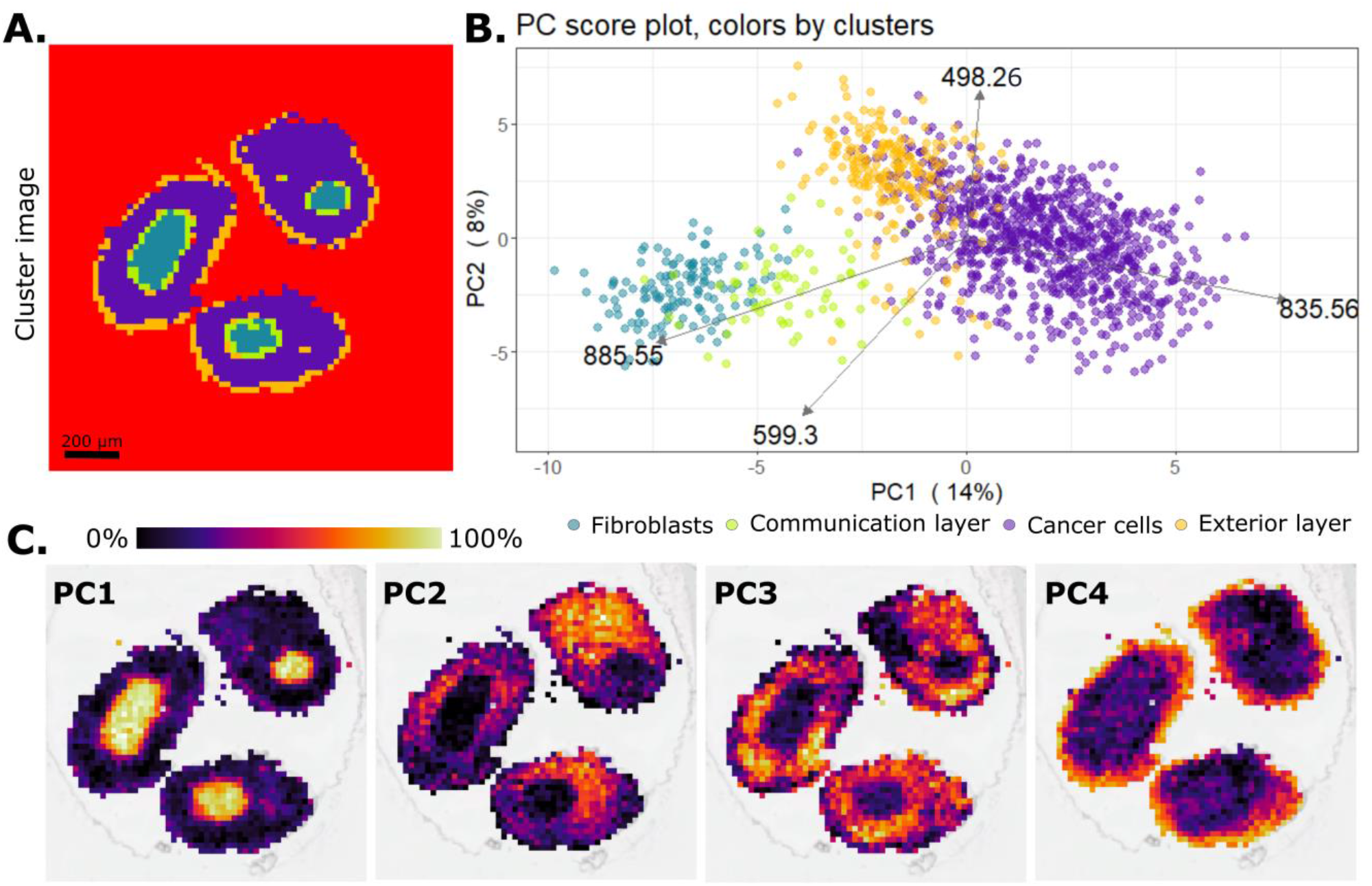
Spheroid MSI data analysis. Segmentation image obtained using bisecting *k*-means clustering (correlation distance) in SCiLS (A); Score plot of PC1 vs. PC2, illustrating the minimum and maximum loadings, pixels colored by regions (B); possible annotations: *m/z* 498.26 as [M-H]^-^ adduct of LysoPE(18:2(9Z,12Z)/0:0), *m/z* 599.3 as [M-H]^-^ adduct of LysoPI(0:0/18:0), *m/z* 835.56 as [M+Na-2H]^-^ adduct of SM(d19:1/PGD1) and *m/z* 885.55 as [M-H]^-^ adduct of PI 38:4; The first four principal component images (C).

We investigated which features are significant in each region by systematic two-by-two volcano plot analysis where the spectral information of all the pixels of one region (cluster) was compared to all the other three regions. Figure 6A shows the volcano plot results of all the possible unique combinations. Significant values were selected by setting the adjusted p value threshold to 0.01 and the fold change threshold to 0.5. We observed that the communication layer is represented by a single significant feature, i.e. *m/z* 867.52, which appears from the comparison with the exterior layer (**Figure 6Ai**). Interestingly, the same feature appears to be significant for the fibroblasts region as well, when compared also to the exterior layer (**Figure 6Aii**). This suggests that finding a feature exclusive to the communication layer is ambitious. This uncertainty could be explained by the molecular similarities of the cells from the communication region and the neighboring cells, grouped in the fibroblast and cancer cell regions. Nevertheless, we found three significant values for the exterior layer, seven for the fibroblasts and four for the cancer cells (**Figure 6B**). The features that have appeared significant for each region in two volcano plot analyses are represented as ion images in **Figure 6C**.

**Figure 6.**
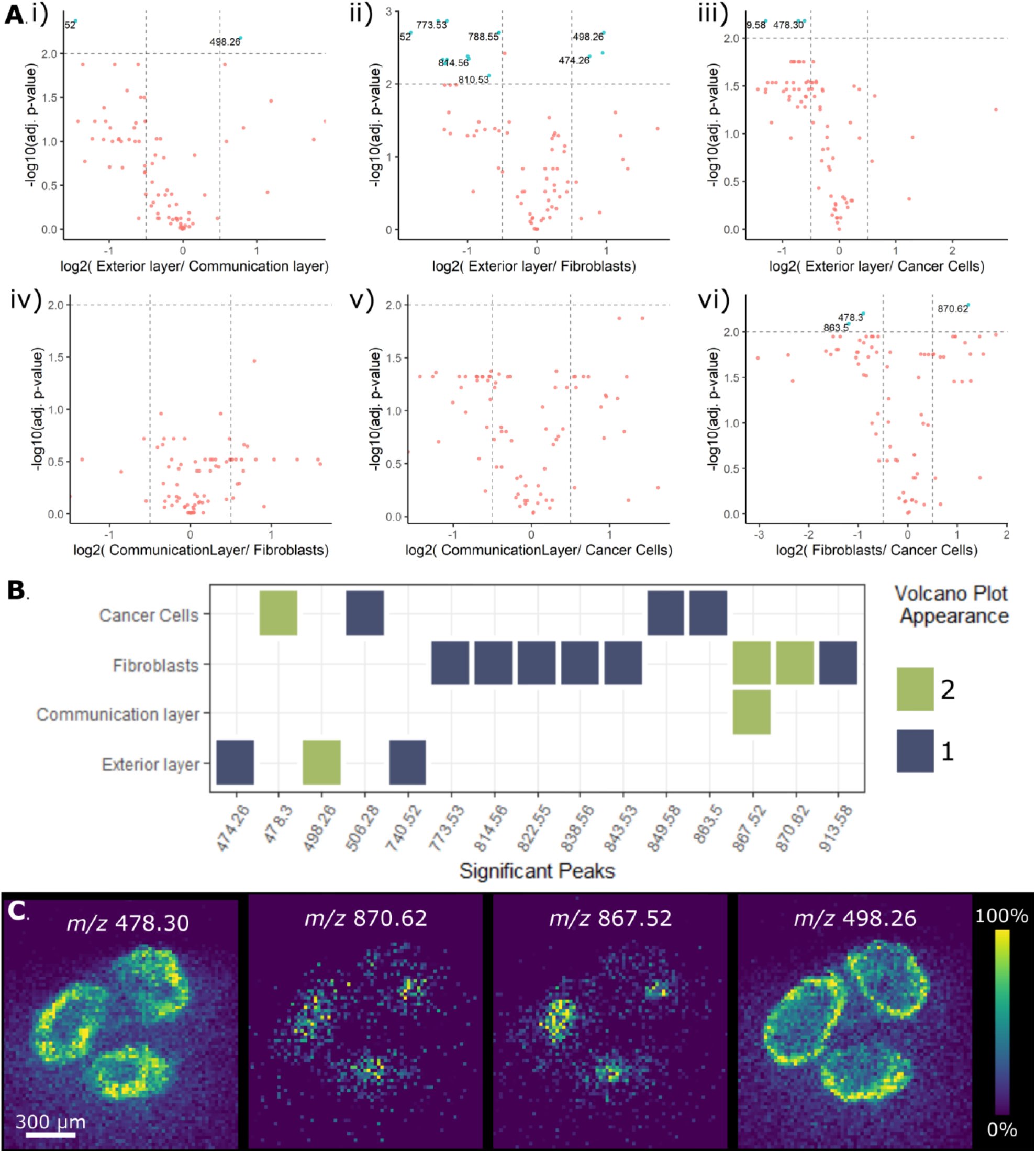
Significant spheroid feature selection. Two-by-two cluster region comparison using volcano plots (A); heat map of significant features for each region showing the number of times the feature has appeared in a volcano plot analysis (B). For volcano plots the adjusted p value threshold was set to padj < 0.01 and the fold change threshold was set to 0.5. Significant ion images representative of each region (C); possible annotations: LysoPE(18:1(11Z)/0:0) for *m/z* 478.3, LysoPE(18:2(9Z,12Z)/0:0) for *m/z* 498.26, PI(16:0/18:1(12Z)-2OH(9,10)) for *m/z* 867.52, PC(20:0/20:3(8Z,11Z,14Z)-2OH(5,6)) for *m/z* 870.62, all presumed to be [M-H]^-^ adducts.

For small molecules, it was possible to generate only two segments within the spheroid regions. These clusters are presumably associated to cancer cells and fibroblasts (**Figure S4A**). In this case, volcano plot analysis was done comparing the two clusters. We set the adjusted p-value and fold change thresholds to 0.05 and 0.5, respectively, and obtained features significant for each cell type (**Figure S4B**). The most significant ion images from each cluster are illustrated in **Figure S4 B** and **C**. Detection in the lower mass range (<300 Da) is challenging, as small molecule signal intensity is generally low and also suppressed by the strong neighboring matrix signals. Therefore, the lower number of spheroid specific features (40 features) – selected using ROC analysis by comparing the matrix and spheroid pixels – could explain why the segmentation was limited to two clusters and why the number of cell-specific significant features is two for each region (or cell type). For this reason, we focused our spatially-oriented data acquisition and analysis in the higher mass range where the two cell lines can be differentiated into multiple metabolically relevant regions.

One of the advantages of 3D cell culture models is their adjustable size. Our bi-cultured spheroids had an average diameter around ~500 μm which allows for the mounting of over 40 spheroid sections on the same ITO slide. Sample size is generally also important for measurement time, which in our case was ~ 5 minutes per section. Our setup used two channels of the gelatin cryo-mold where one position was filled with bi-cultured spheroids, with 45 consecutive sections on one Intellislide (**Figure 7A**). Thus, it was possible to collect data in the range of *m/z* 300-1200 for 3D reconstruction in about 6 hours.

**Figure 7.**
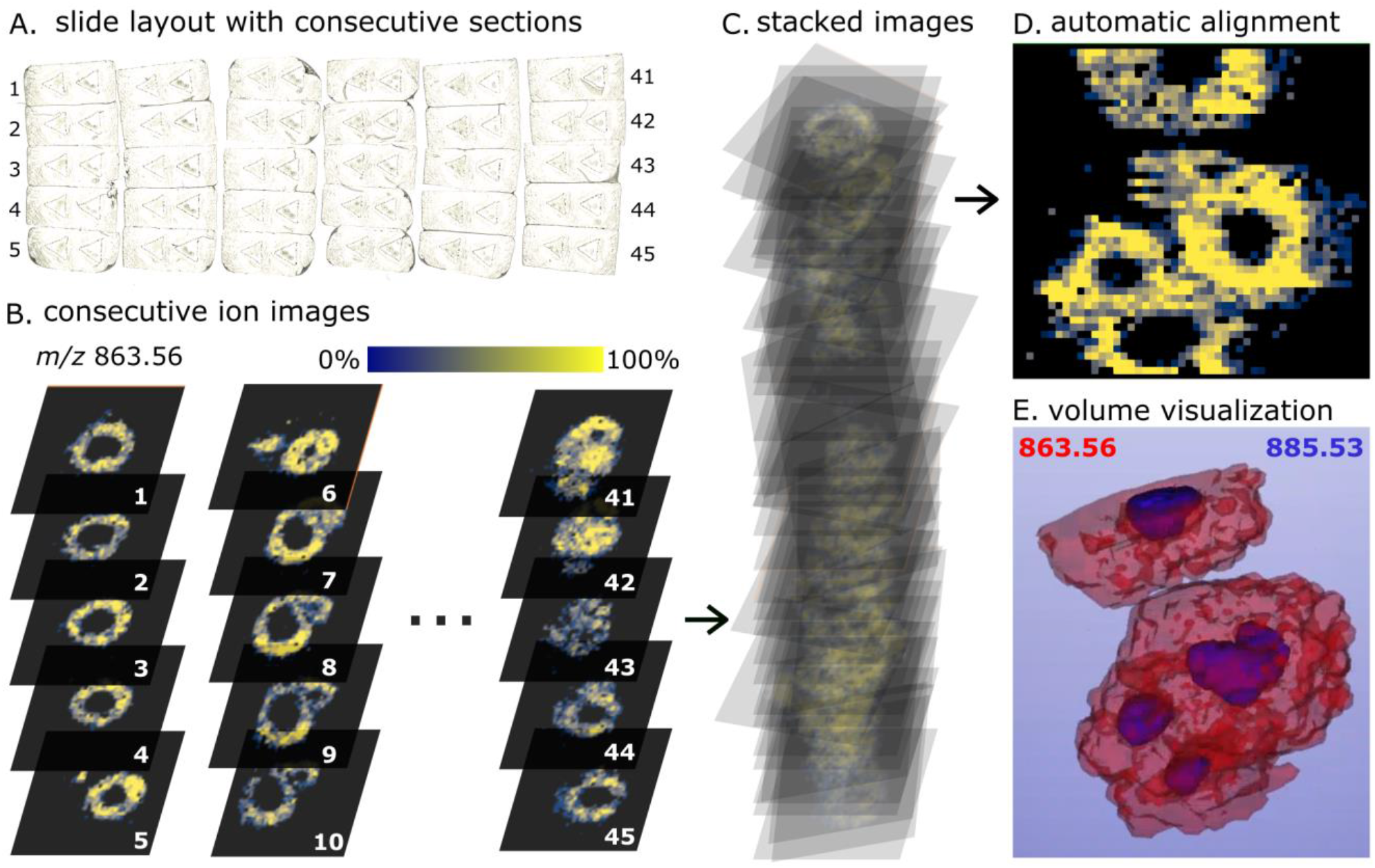
3D reconstruction strategy. Slide layout with consecutive sections (A); Consecutive sections illustrated by the ion images of *m/z* 863.56 (B); automatic stacking and registration of consecutive ion images using M^2^aia software: illustrative stacking (C) and cross-sectional view of stack (D); volume visualization of different ions of interest, in red *m/z* 863.56 and in blue *m/z* 885.53 (E).

For fast visualization of specific ions from all sections, the data was then imported into SCiLSlab software. This was followed by export of imzML files of each section individually and their import into M^2^aia software[19] for automated 3D reconstruction (**Figure 7BC**). *m/z* 863.56 was selected as landmark for registration in creating the stacked image (**Figure 7D**), from which the stacked images of region-specific features were then extracted: *m/z* 498.26 for the exterior layer, *m/z* 835.56 for the cancer cells and *m/z* 885.55 for the fibroblasts. Then, the stacked images were visualized in 3D using Slicer 3D software (**Figure 7E**).

**Figure 8.**
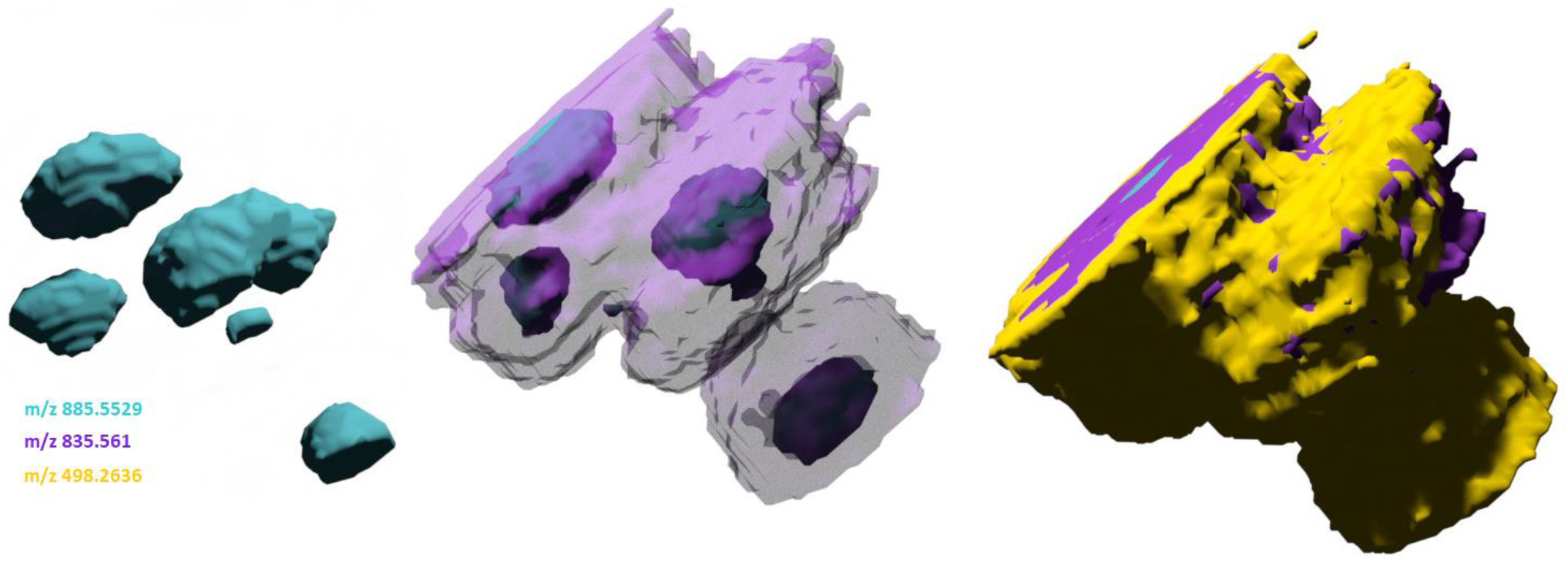
Representation of region-specific ion distributions in space. Frames from a 3D reconstruction video (Supporting file 1) with *m/z* 498.26, *m/z* 835.56, and *m/z* 885.55 describing the exterior layer (yellow), the cancer cells (purple), and the fibroblasts (cyan).

Previously, biologically relevant regions determined by *k*-means clustering were shown in 3D[15], hinting to a realistic interpretation of different phenotypical features of the tissue. In doing so, Flint et al. were able to separate and identify metabolites regulating cancer growth from distinct regions, thus mimicking an important aspect of the heterogeneity of *in vivo* tumors. Similarly, we have shown that the molecular composition of co-cultured spheroids can be used to identify four metabolically distinct zones in bi-culture spheroids that suggest interactions between the two cell types, which promote the formation of additional metabolically distinct zones such as the “communication” layer. Here, we were able to clearly distinguish the two cell types of the co-culture (fibroblasts and cancer cells) by finding discriminating features of each cell type. Further work is necessary to unequivocally assign the different zones to functionally relevant cellular functions, such as specific cellular signaling or metabolic interaction between cancer and fibroblast cells. Multi-modal imaging combining immunohistochemical staining and MSI could shed light on the morphological but also functional aspects of this region, while LC-MS/MS coupled to MSI through laser capture microdissection (LMD)[23] could provide a better overview of the chemical composition of these cells.

## Conclusion

In this work, we presented a platform for MALDI MSI of 3D cell culture models, which enables rapid and reliable preparation of man serial sections of spheroids on a single ITO slide, followed by spatial metabolomic profiling and elastic 3D reconstructions of molecular patterns. We demonstrated that cell type and cell layer-specific metabolites can de distinguished by multivariate analysis, suggesting the possibility to identify relevant markers for biological mechanisms typical to each cell type.

## Acknowledgments

The authors would like to acknowledge the funding by the German Federal Ministry of Education and Research (BMBF) as part of the Innovation Partnership M^2^Aind, projects ADCtox-3D (***13FH8E01IA***) and Drugs4Future (***13FH8I05IA***) and M^2^Aind-DeepLearning (***13FH8I08IA***). JC and IW acknowledge the funding from Digi-FIT project within the framework FH-Impuls and by the Carl-Zeiss-Foundation. The authors also acknowledge the help from Mathias Rädle and Lucas Schmitt for improving and manufacturing the metal casting mold and engaging discussions about 3D cell culture models with Elina Nurnberg and Ieva Palubeckaite.

