## Supplementary material for "3D-Mass Spectrometry Imaging of Micro-scale 3D Cell Culture Models in Cancer Research": Iakab_etal_SupportingInformation.pdf

**Affiliations:**

**Contents:**

Table S1. Spraying protocols used on the HTX M3 sprayer

Figure S1. 3D-printed metal casting molds for nine and three channels

Figure S2. Spheroid embedding workflow

Figure S3. Consecutive sectioning and mounting on ITO slide

Figure S4. Spheroid MSI data analysis for small molecules

**Table S1. Spraying protocols used on the HTX M3 sprayer**. Constant parameters: 1200 mm/min velocity; 2 mm track spacing; CC pattern; 10 psi pressure; 0 s dry time; 40 mm nozzle height.

| Matrix | Concentration (mg/ml) | Solvent | Temp (°C) | Number of passes | Flow rate (µl/min) |
|---|---|---|---|---|---|
| 9-AA | 6 | 70%ACN | 60 | 10 | 0.08 |
| DAN | 6 | 70% ACN | 60 | 10 | 0.06 |
| DHB | 30 | 70% ACN | 60 | 15 | 0.05 |
| NEDC | 10 | 70% methanol | 55 | 8 | 0.08 |
| PhCCAA | 3 | 70% ACN | 60 | 12 | 0.08 |
| BNDM | 10 | 70% ACN | 55 | 16 | 0.07 |

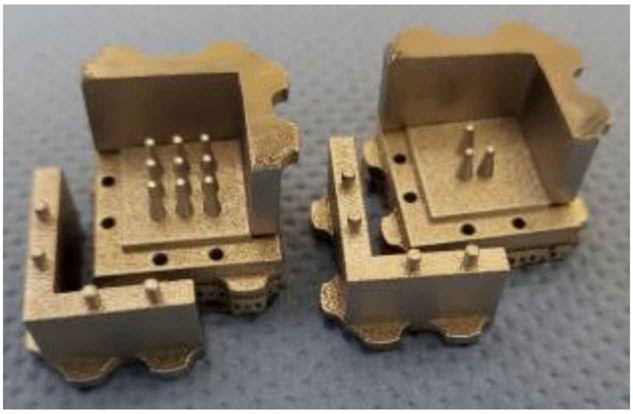

**Figure S1**. **3D-printed metal casting molds for nine and three channels**

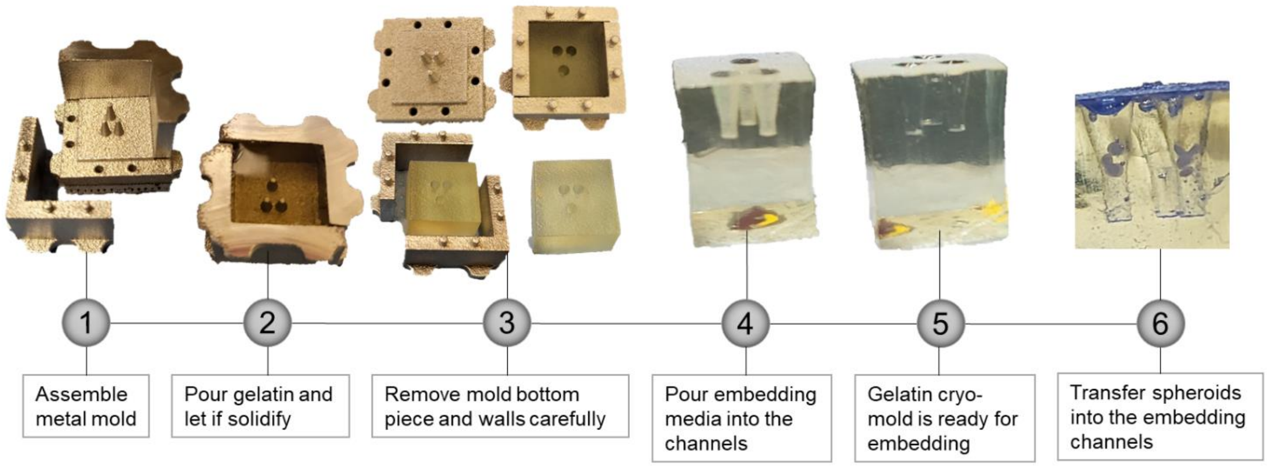


**Figure S2. Spheroid embedding workflow.** Schematic step-by-step illustration of the embedding process (photos ©CeMOS).

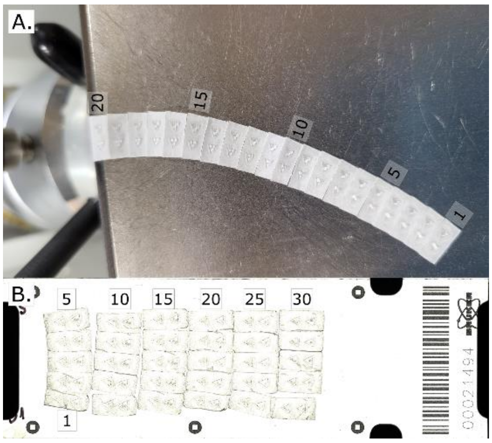


**Figure S3. Consecutive sectioning and mounting on ITO slide**. Twenty consecutive gelatin cryosections were obtained (A) and 30 consecutive sections were mounted on an Intellislide (B).

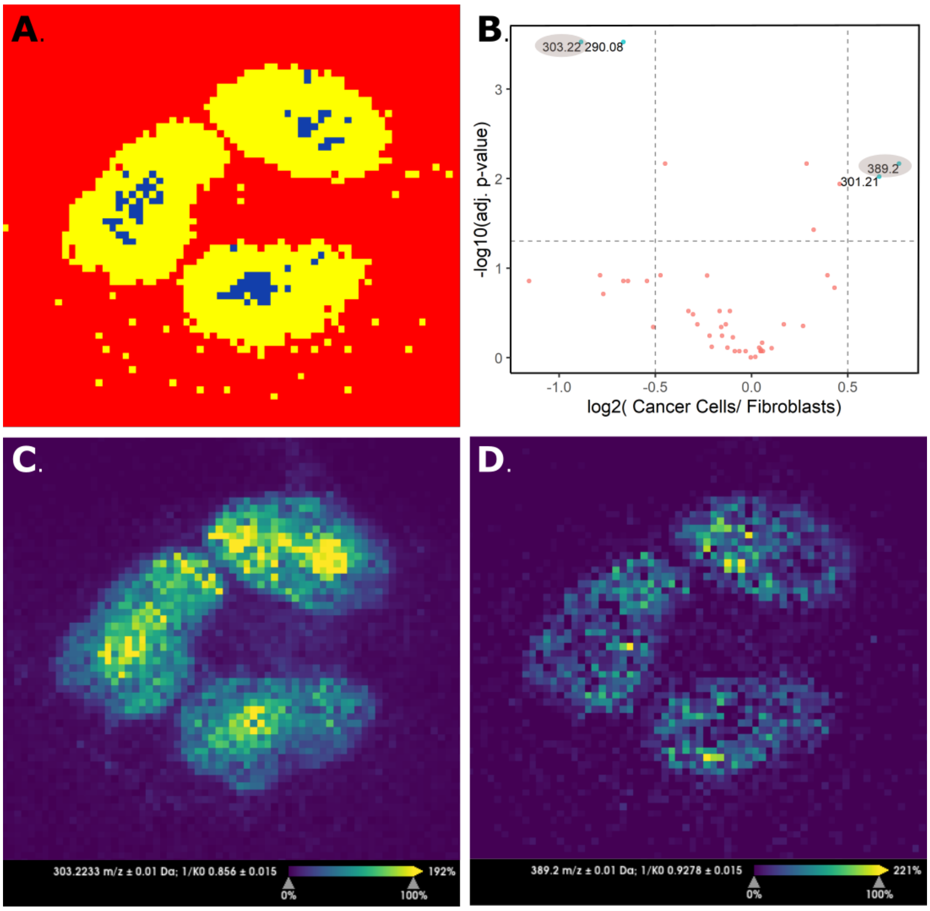


**Figure S4. Spheroid MSI data analysis for small molecules**. Segmentation image obtained using bisecting k-means clustering (correlation distance) in SCiLS (A); Volcano plot comparing cancer cells cluster (in yellow) and fibroblast cluster (in blue) (B); Ion images of the most significant feature of fibroblasts (C) and cancer cells (D).
